# Single-cell analysis reveals the cellular and transcriptional diversity of thyrocytes in the normal pediatric thyroid

**DOI:** 10.1101/2025.11.17.687863

**Authors:** Erin R. Reichenberger, Nicholas E. Bambach, Zachary Spangler, Julio C. Ricarte-Filho, Kyle Hinkle, Amber Isaza, Tricia Bhatti, Andrew J. Bauer, Aime T. Franco

**Affiliations:** Department of Biomedical and Health Informatics, Children’s Hospital of Philadelphia, Philadelphia, PA, United States; Division of Endocrinology and Diabetes, Children’s Hospital of Philadelphia, Philadelphia, PA, United States; Department of Pathology and Laboratory Medicine, Children’s Hospital of Philadelphia, University of Pennsylvania, Philadelphia, PA, United States; Abramson Cancer Center, University of Pennsylvania, Philadelphia, PA, United States

**Keywords:** pediatric, thyroid, scRNA-seq, thyrocytes, transcriptional, heterogeneity, hormone, biogenesis

## Abstract

To enhance the understanding of cellular heterogeneity within the pediatric thyroid, single-nuclei RNA sequencing was used to recover 38,069 non-pathogenic cells from thyroid tissue of three pediatric patients. The recovered cells were analyzed using the SWANS (**S**ingle Entity **W**orkflow **AN**alysi**S)** pipeline (version 1.0). Analysis revealed seven major cell types: thyrocytes, endothelial cells, fibroblasts, C cells, T cells, B cells, and myeloid cells. Thyrocytes were the most prominent and heterogeneous cell type. Initially, two dominant thyrocyte subsets were identified based on transcriptional activity, which were subsequently subdivided into seven subclusters. Differentially expressed genes within each cluster support distinct cellular functions, including a metabolically active subset which may be involved in hormone synthesis and a subset involved in the transport of thyroid hormone into circulation. We identified an immune subpopulation originating predominantly from a single sample that was histologically and morphologically similar to the other two samples. This supports that transcriptional changes can be detected and used to identify populations of cells, even in the absence of histologically observable changes. This characterization represents the first comprehensive portraiture of pediatric thyroid gland cells and the first description of normal patient thyrocyte and stromal cell heterogeneity in the absence of adjacent malignancy.

## INTRODUCTION

The thyroid is a bilateral symmetric endocrine organ located in the anterior lower region of the neck (Figure 1A) which primarily produces the hormones thyroxine (T4) and triiodothyronine (T3). Thyroid hormones play a central role in metabolism, growth, and development, and often serve as a rheostat between multiple endocrine systems to coordinate physiologic homeostasis. Thyroid dysfunction is quite common, affecting 100 million people worldwide (1). Hypothyroidism can be managed successfully with thyroid hormone replacement, while hyperthyroidism is often managed effectively through medication that inhibits thyroid hormone production, thyroid ablation, or surgery. When left untreated, either condition can result in lifelong morbidities and decreased quality of life, particularly in children (2).

**Figure 1.**
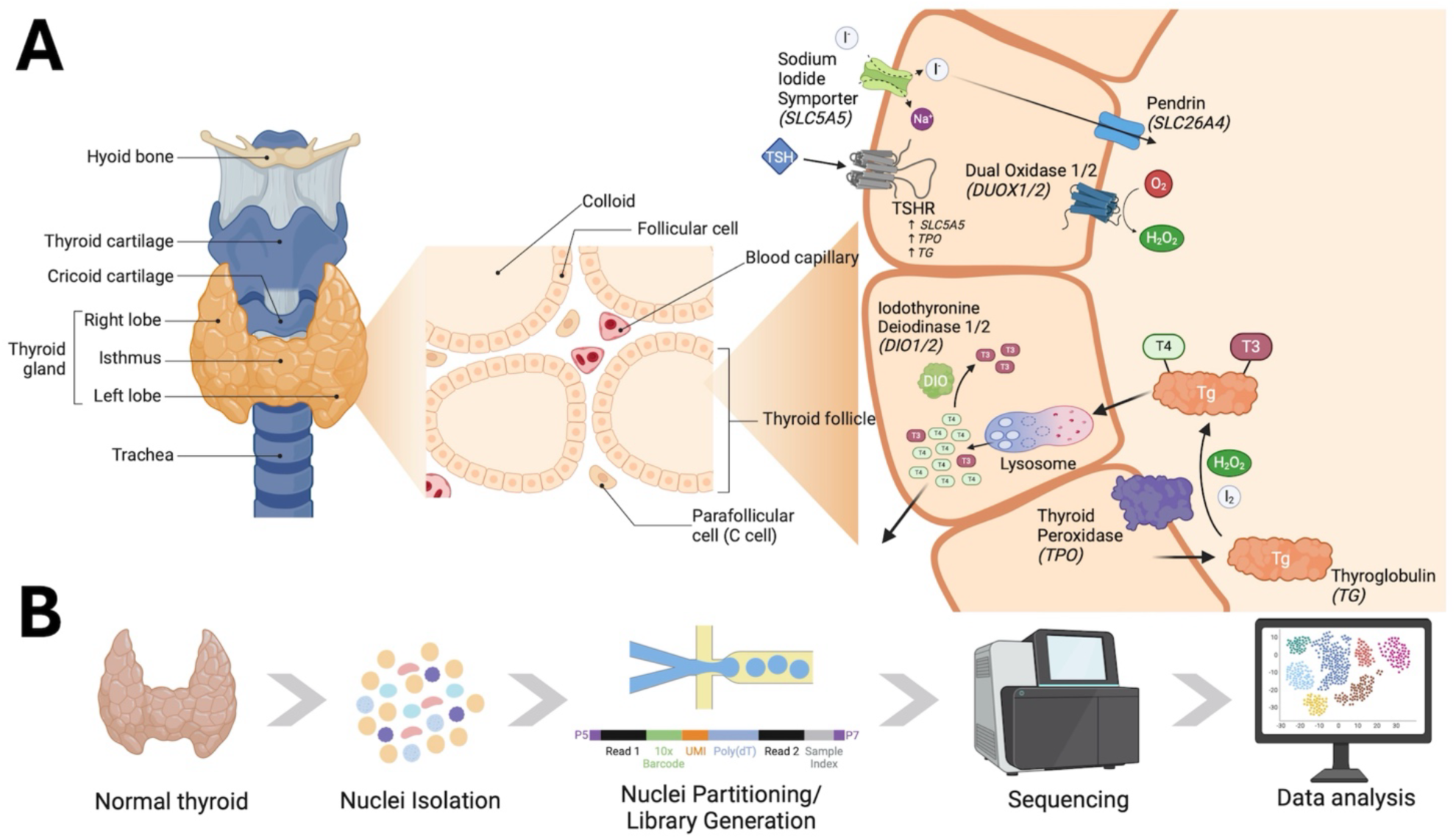
Thyroid gland physiology and study design. **(A)** Image depicting the anatomical location of the thyroid gland, zooming in to illustrative histology and further depicting the cellular localization of key proteins involved in thyroid hormone biogenesis. **(B)** Schema representing study design and experimental processing of thyroid tissue for snRNA sequencing. Both images were created and adapted using BioRender.com.

Production, storage, and secretion of thyroid hormones by thyrocytes is tightly regulated through the hypothalamic-pituitary-thyroid axis. Thyrotropin releasing hormone is secreted from the hypothalamus, which stimulates the production and release of thyroid stimulating hormone (TSH) from pituitary thyrotrophs. TSH binds to the TSH receptor (TSHR) on thyrocytes and stimulates the synthesis and secretion of thyroid hormones, in addition to acting as a mitogenic stimulus to thyrocytes. The thyroid itself is organized into follicles of various sizes, which produce and store hormones for on-demand release in response to TSH. Thyroid hormone synthesis begins with iodine transport into thyrocytes mediated by the sodium iodide symporter (NIS) encoded by *SCL5A5*. Iodine is then coupled to tyrosine residues on thyroglobulin (TG), a reaction which is catalyzed by thyroid peroxidase (TPO). Iodinated TG is stored within the colloid of the thyroid follicles which can then be transported back through thyrocytes, cleaved by cysteine proteases within lysosomes to form T3 and T4, and released into neighboring blood vessels for transport to distant sites within the body (Figure 1A). Although the biochemistry of thyroid hormone biosynthesis has been well-defined, it remains unknown whether all thyrocytes in the human thyroid have the same ability and functionality to produce and secrete thyroid hormones. Gillotay and colleagues reported thyrocyte heterogeneity within the zebrafish, but definition of such phenomena within the normal human gland, particularly within the pediatric population, remains incomplete (3).

The thyroid gland is comprised of many different cell types. This includes a small subset of thyroid cells called C cells or parafollicular cells, which are responsible for calcitonin production. The thyroid is a highly vascularized organ which uptakes iodine for thyroid hormone biosynthesis and transports thyroid hormone throughout the body. In thyroid cancer, robust immune recruitment is often observed, as well as fibroblast-mediated remodeling of the extracellular matrix (4–9). In the pathological conditions of Graves’ disease and Hashimoto’s thyroiditis, immune remodeling and expansion of stromal cells are also observed. However, the presence and role of immune cells and fibroblasts in the normal thyroid have not been well characterized or defined. Much of our knowledge of the cell types present in the normal thyroid has been defined through work in model organisms or in the context of thyroid malignancy (3, 10–12). Limited access to normal thyroid tissue in the absence of adjacent malignancy has hindered our ability to define the cellular landscape of the normal thyroid gland. Furthermore, much of our understanding of the physiology of the normal thyroid was generated before tools such as single-cell RNA sequencing were available. Thus, we lack molecular characterization of the diversity that exists within thyrocyte populations and the multiple cell types that are likely present within the normal pediatric thyroid gland. Here, for the first time, we profile cells from pediatric thyroid tissue, collected from thyroids with no background malignancy.

Recent advances in single-cell and single-nucleus RNA sequencing (scnRNA-seq) have enabled high-resolution transcriptional profiling at the level of individual cells, offering critical insights into cellular heterogeneity, the identification of rare cell types, lineage tracing, and the elucidation of complex biological processes (13–17). While there are numerous computational tools and pipelines developed to facilitate the processing and analysis of scnRNA-seq data (18–26), we took advantage of the flexibility and highly-customizable framework provided by SWANS (27) to define multiple cell types within the pediatric thyroid gland from three female patients with no overt pathological changes. This study presents the first single-cell transcriptional characterization of the human thyroid gland using samples derived entirely from non-malignant pediatric patients. This data set has confirmed the presence of heterogeneous populations of thyrocytes, resident fibroblasts, endothelial cells, parafollicular cells, and immune cells within the pediatric thyroid gland, and will be a valuable reference for future studies.

## MATERIALS AND METHODS

### Patient selection and approval

Three age and sex matched pediatric patients who had undergone partial thyroidectomy for suspicious nodules which were subsequently diagnosed as benign thyroid conditions were selected for this study (Table 1). Each patient had a confirmed benign diagnosis after surgical removal and pathological inspection of the nodule. Adjacent normal tissue that had been flash frozen at the time of surgery was used for this study and single nuclei were isolated and prepared as described below.

**Table 1.**
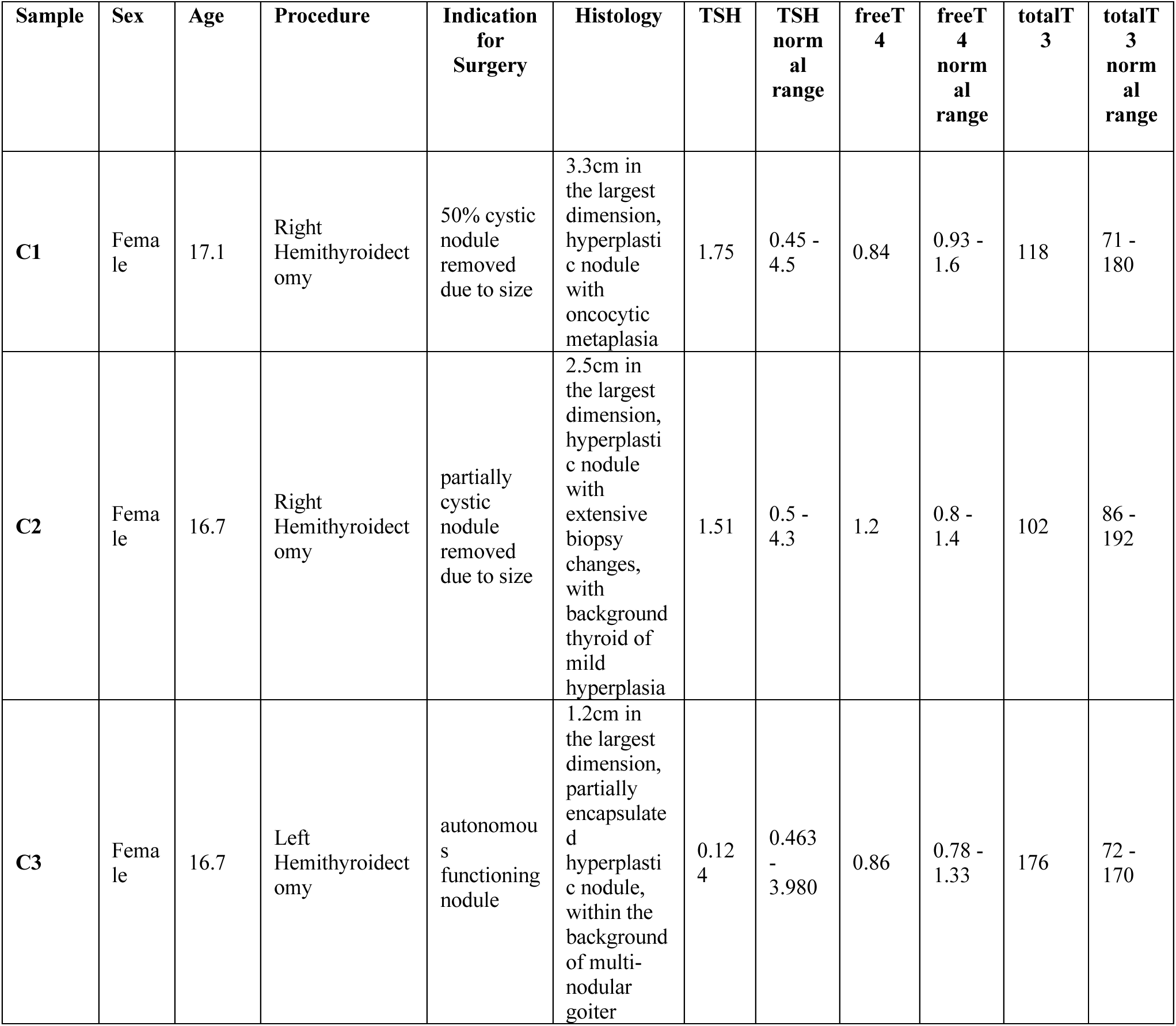
Patient demographics and clinical diagnosis for samples used for snRNA sequencing analysis.

### Tissue dissociation and single nuclei preparation

Frozen non-neoplastic thyroid samples (∼30-40 mg, n=3) were dissociated into nuclei suspensions using the Chromium Nuclei Isolation Kit (10x Genomics) according to manufacturer’s instructions. Thyroid tissues were dissociated with a pestle in 200 µl of ice-cold lysis buffer until homogeneous. Next, 300 µl of lysis buffer were added to the samples and then incubated on ice for 10 minutes. Samples were then transferred to a pre-chilled Nuclei Isolation Column and centrifuged at 16,000 rcf for 20 seconds at 4°C. Flowthrough was vortexed at 3,200 rpm to resuspend nuclei and centrifuged for 3 minutes at 500 rcf at 4°C. Nuclei pellets were resuspended in 500 µl of Debris Removal Buffer and centrifuged at 700 rcf for 10 min at 4°C to remove debris. The isolated nuclei were then washed twice with Wash and Resuspension buffer containing RNase inhibitor and centrifuged at 500 rcf for 5 min at 4°C. The nuclei suspension was adjusted with Wash and Resuspension buffer (containing RNase inhibitor) to a density of 800-1,000 cells/μl to yield 16,500 cells.

### Single nuclei sequencing of the pediatric thyroid gland

The nuclei samples were sequenced at the CHOP Single Cell Technology core facility. Nuclei were mixed with the reverse transcription (RT) mix and partitioned into GEMs (Gel Beads-in-EMulsion) using the Chromium Controller. To facilitate the RT reaction, one hundred microliters of recovered GEMs were incubated at 53°C for 45 minutes, then 85°C for 5 minutes, followed by a 4°C hold. After the RT reaction, GEMs were broken, and the first stand of cDNA product was collected and cleaned using Dynabeads MyOne SILANE magnetic beads. The cDNA was then amplified for 11 cycles using the protocol outlined in the referenced user guide. The 3’ Gene Expression Library preparation involved cDNA fragmentation, end repair and a-tailing, adaptor ligation, and sample index PCR. Library samples were indexed using individual sample index sets (Dual Index Plate TT Set A). The number of PCR cycles was determined based on the cDNA input measured by Bioanalyzer. Thirteen cycles of amplification were performed for all samples. Indexed samples were then cleaned up using a 0.6-0.8X SPRIselect magnetic bead double size selection. 35 μl of purified product were used, and library concentration was measured by Qubit dsDNA HS Assay Kit (Invitrogen Q32851). The libraires were sequenced on Illumina’s NovaSeq 6000 S4 v1.5 flow cell for 26x10x10x90 cycles to obtain a sequencing depth of 30,000 reads/nucleus. FASTQ files were aligned to the GRCh38 human reference genome using 10X Genomics’ Cell Ranger (version 7.1.0) ‘count’ pipeline with ‘--include-introns’ and default parameters (26). Some of the resulting files (*e.g*., features, matrix, barcodes) represent the starting data for the analysis.

### Preprocessing and analysis of single nuclei sequencing

Using the SWANS pipeline and the output from the Cell Ranger’s ‘count’ pipeline, per-sample ambient mRNA contamination was evaluated and removed using SoupX (version 1.6.2) (28). Multi-cellular droplets were identified and removed using DoubletFinder (version 2.0.3) (29). Per-sample Seurat (version 4.3) objects were created and collectively merged into a single Seurat object for subsequent analysis (17, 19). Cells expressing less than 200 or more than 3,000 genes were excluded from the object. Cells with more than 15% of features attributed to mitochondrial genes were also excluded. Expression data was then normalized using Seurat’s “NormalizeData” function. The top 2000 most variable genes were found (“FindVariableFeatures”) and scaled for use in downstream analysis. Initially, the Seurat object was analyzed using 50 principal components (PCs) and the amount of variance in each component was calculated. The number of components which collectively contributed 90% of the variance (12 PCs) was retained for analysis. Thyroid tissue cells were analyzed following Seurat’s RPCA approach at multiple resolutions (0.1-0.8 by 0.1). UMAP figures were created for each resolution and upregulated differentially expressed genes (DEGs) were identified for all clusters at each resolution using Seurat’s ‘FindAllMarkers’ command with default parameters. Results for all available clustering arrangements were inspected with SWANS’ interactive report. To assist with schema selection, the Clustree package (version 0.4.3) was used, and the final clustering schema was based off a combination of multiple resolutions that allowed for the highest number of clusters while obviating any over clustering (30).

### Identification of specific cell types within the thyroid, sample similarity, and pathway analysis

To identify annotation markers unique to and strongly expressed in a single cluster, the DEGs expressed in at least 25% of cells within a given group with a minimum average log-fold-change of 4.0, were selected using Seurat’s ‘FindAllMarkers’ command with default parameters. When necessary, individual clusters were combined into one cluster and Seurat’s ‘FindMarkers’ command was employed again using default parameters on the combined cluster against all remaining clusters to find upregulated genes. To identify C cells, the number of *CALCA* ‘counts’ per cell was obtained from the gene count matrix. For each sample, cells with zero *CALCA* counts were removed, and the remaining values were converted to Z-scores. Cells with Z-scores ≥ 1.96 (∼ confidence interval 95%, p-value ≤ 0.05) were noted and used with the ‘FindMarkers’ function to find upregulated DEGs. A UMAP image of the final clustering schema was made, and the number and proportion of cells in each cluster were calculated for each sample. To assess gene expression similarity across samples, the average expression of each gene (per sample) was calculated with Seurat and the correlation between paired samples was calculated with the ‘cor.test’ function in the corrplot R package (version 0.92) (31). Seurat’s ‘FindConservedMarkers’ was used to identify genes with similar expression patterns across samples. The resulting gene list was ranked by average log2 fold change and analyzed using the fgsea R package with the Molecular Signatures Database (MSigDB) to identify enriched biological pathways (32–35). Pathways were restricted to those from the hallmark (H), curated (C2), and ontology (C5) collections. Results were filtered for an adjusted p-value of 0.1 and were further filtered for entries by investigators for processes of interest related to normal thyroid function.

### Subclustering of thyrocytes

Cells belonging to thyrocyte populations in each sample were separated from other cell types. These cells were analyzed with the SWANS pipeline using similar methods as described above. The Seurat object was initially analyzed with 50 PCs and the amount of variance per PC was calculated. The number of components (13 PCs) that captured 90% of the variance was noted and used to analyze the thyrocyte cells using Seurat’s RPCA approach at multiple resolutions 0.1 - 0.8 (by 0.1). UMAP images of each clustering arrangement were created for each resolution, and the DEGs were examined in SWANS’ interactive environment. Clustree was again employed to identify the optimal number of clusters while avoiding over-clustering, and multiple resolutions were integrated into the final clustering framework. Violin plots of QC metrics for each cluster were made with Seurat. To identify biological processes affiliated with each thyrocyte subcluster, genes were ranked based on their Z-transformed expression value. These ranked lists were used for pathway analysis with the fgsea package and the Molecular Signatures database (MSigDB) (hallmark, curated, and ontology gene sets). Results were filtered for an adjusted p-value of 0.1. Pathways shared across all clusters and pathways unique to a cluster were found using ‘intersection’ and ‘union’ mathematical methods. To identify genes uniquely expressed in a thyrocyte subcluster, we first averaged the expression of each gene across cells within each cluster. For each cluster, we then determined which genes had an average expression that was at least three-fold higher in the given cluster compared to all other clusters. Genes with low expression across all clusters were not evaluated.

## RESULTS

### Understanding the cellular heterogeneity of the pediatric thyroid gland

To investigate the cellular heterogeneity of the pediatric thyroid gland and the associated molecular signatures of the observed cell types, we profiled thyroid tissue from three age-matched female pediatric patients (Figure 1B, Table 1). All tissue was collected from adjacent normal tissue of patients undergoing lobectomy for suspicious nodules and was flash frozen at the time of diagnosis. Each nodule was diagnosed as benign post-operatively following histological inspection by an experienced pediatric pathologist (Figure S1).

After QC and processing of the snRNA-seq data with SWANS, a total of 36,977 cells were analyzed, with the number of recovered cells per sample ranging from 10,869 to 13,746 (Table 2). To date, this represents the largest number of cells profiled from a single thyroid sample, making this the most comprehensive thyroid tissue data set – pediatric or adult. A summary of cellular features, RNA fragments, and mitochondrial percentages can be seen in Table 3.

**Table 2:** Table of cells (and their proportions) for each of the 9 clusters identified in three pediatric thyroid glands.

|  | **C1** | **C2** | **C3** | **Total** |
| --- | --- | --- | --- | --- |
| **Thyrocyte0** | 9361 (68.1) | 8150 (65.93) | 6952 (63.96) | 24463 (66.16) |
| **Thyrocyte1** | 3375 (24.55) | 2759 (22.32) | 3405 (31.33) | 9539 (25.8) |
| **Endothelial_L** | 366 (2.66) | 525 (4.25) | 196 (1.8) | 1087 (2.94) |
| **Endothelial_V** | 209 (1.52) | 57 (0.46) | 79 (0.73) | 345 (0.93) |
| **Fibroblasts** | 269 (1.96) | 154 (1.25) | 134 (1.23) | 557 (1.51) |
| **Mixed_Tcells** | 98 (0.71) | 477 (3.86) | 62 (0.57) | 637 (1.72) |
| **Myeloid** | 54 (0.39) | 75 (0.61) | 41 (0.38) | 170 (0.46) |
| **C.Cells** | 13 (0.09) | 2 (0.02) | 0 (0) | 15 (0.04) |
| **Mixed_Bcells** | 1 (0.01) | 163 (1.32) | 0 (0) | 164 (0.44) |
| **Total** | 13746 (100) | 12362 (100) | 10869 (100) | 36977 (100) |

**Table 3:** Summary statistics (number of genes and molecules per cell, percentage of mitochondrial genes per cell) of datasets.

|  | **nCount_RNA** | **nFeature_RNA** | **percent.mito** |
| --- | --- | --- | --- |
| Min. | 421.23 | 207.00 | 0.00 |
| 1st Qu. | 1650.93 | 1142.00 | 0.00 |
| Median | 2292.14 | 1467.00 | 0.09 |
| Mean | 2468.57 | 1502.07 | 0.75 |
| 3rd Qu. | 3119.24 | 1832.00 | 0.75 |
| Max. | 8160.73 | 2999.00 | 14.86 |

### Cluster annotation and proportions

Nine clusters were initially identified in the dataset. After reviewing the DEGs affiliated with two suspected fibroblasts clusters, it was decided to manually combine the two populations into one cluster. Additionally, as described in the methods, we manually identified C cells based on *CALCA* expression and relabeled these selected cells as C cells. A UMAP denoting the final nine clusters and seven major cell types of the thyroid dataset can be seen in Figure 2A. The markers used to identify the clusters can be visualized in Figure 2B and are arranged by sample. Visual and numerical representations of the number and proportion of cells in each cluster are presented in Figure 2C and Table 2, respectively.

**Figure 2.**
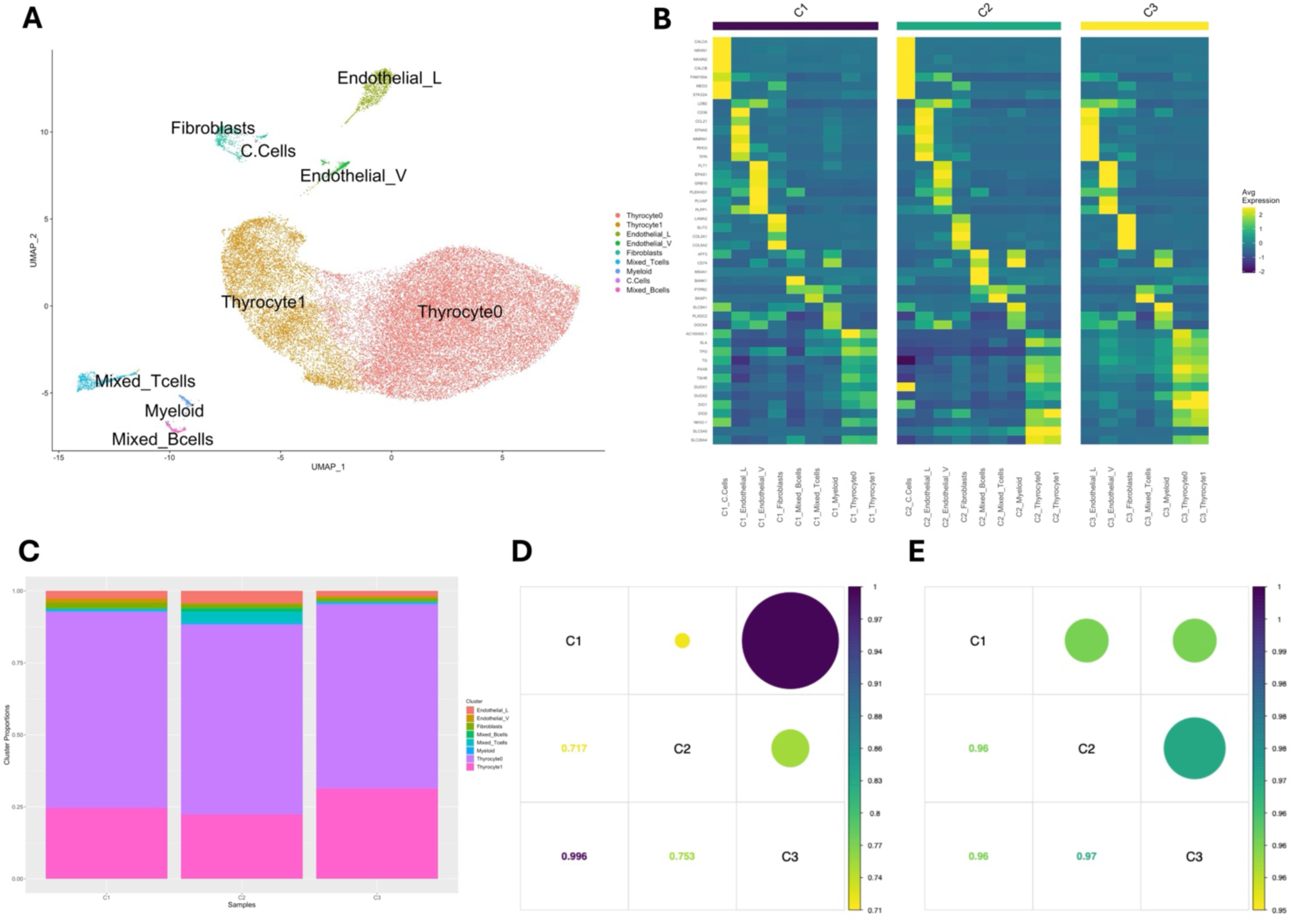
Identification of cell types within the thyroid gland. **(A)** Final UMAP embedding of 36,977 cells derived from three pediatric thyroid glands. Each colored cluster contains transcriptionally similar cells and represents nine cell types. **(B)** Heatmap depicting averaged normalized expression of the markers used for cluster annotation. For non-thyrocytes, an adjusted p-value <= 0.05 and an average log2FC >= 4.0 were the selection criteria for annotation makers found with Seurat’s ‘FindAllMarkers’ function. Thyroid differentiation score markers were used to annotate thyrocytes. **(C)** Bar plot depicting relative abundance of cells within each cluster and independently across three pediatric thyroid glands. **(D)** A correlation plot displaying Spearman’s correlation coefficients based on cluster proportions across the three pediatric thyroid samples. **(E)** A correlation plot displaying Spearman’s correlation coefficients* based on sample transcriptional profiles across the three pediatric thyroid samples. *Correlations were run on the average normalized expression of all genes in dataset.

Thyrocytes (clusters Thyrocyte0 and Thyrocyte1) were identified by canonical thyroid genes *TPO, TG, TSHR, PAX8, DUOX2, DIO1, DIO2, NKX2-1*, and *SLC26A4*. These clusters represented the most abundant cell type and constituted almost 92% of all cells in the dataset. Endothelial cells were the next most abundant cell type, comprising 3.9% of all profiled cells. Two distinct endothelial clusters were identified: vascular (∼3%) and lymphatic (∼0.9%) endothelial cells. Vascular endothelial annotation markers included *FLT1, EPAS1, GRB10, PLEKHG1, PLVAP*, and *PLPP1*. *FLT1* is a member of the vascular endothelial growth factor receptor (*VEGFR*) family and plays a role in angiogenesis and vasculogenesis (36). For lymphatic endothelial cells, *LDB2*, *CD36, CCL21, EFNA5, MMRN1, RHOJ*, and *TFPI* were the most prominently expressed markers. The marker *CCL21* is a chemokine particularly important for lymphangiogenesis and can be upregulated in response to inflammation (37, 38). Both sets of markers have previously been associated with vascular and lymphatic endothelial cells.

Fibroblasts accounted for 1.5% of cells and were identified based on expression of *LAMA2, SLIT2, COL3A1*, and *COL5A2.* These cells originally clustered into two independent groups, but upon inspection of the markers, were combined into a single fibroblast population. It is possible that there are multiple independent resident fibroblast populations within the thyroid; however, the small number of cells hindered our confidence in concluding whether they represent distinct entities. All identified immune cells expressed *PTPRC* (*CD45*). *SKAP1* was enriched within the mixed T cell population, which represented 1.7% of cells. B cells (∼0.44%) were marked by increased expression of *AFF3, CD74, BANK1, MS4A1*, and *ARHGAP15*, whereas myeloid cells (∼0.46%) had the highest expression of *SLC8A1* and *SRGN*. Initial clustering did not isolate or identify a distinct population of C cells. However, C cells were identified using statistically significant z-score transformations on cellular *CALCA* expression, as described in detail under Materials and Methods.

C cells were only identified in samples C1 and C2, and represented only 0.041% of all profiled cells (Tables 2, S1A). The most highly expressed markers for C cells were *CALCA, NRXN1, NKAIN2, CALCB, FAM155A, MEG3,* and *STK32A*. Genes *CALCA* and *CALCB* are both members of the calcitonin family. Although there are no studies to our knowledge that have profiled normal human C cells, *FAM155A, MEG3,* and *STK32A* were all reported as markers of C cells in murine snRNA-seq analysis in unpublished data from the Franco laboratory. The identified C cells were initially classified as fibroblasts (11 cells), thyrocytes (3 cells), and a vascular endothelial cell (1 cell).

### Sample similarity

Cells from each individual patient sample were present in most of the identified clusters. The most variation between samples’ cell presence and proportion were found in clusters containing very few cells (Figure 2C, Table 2). Trivial differences in the total proportion of thyrocytes were observed between the three samples. The proportion of fibroblasts was consistent across all three samples, while C cell identification was rare and restricted to samples C1 and C2. Lymphatic endothelial cells were more abundant than vascular endothelial cells across all three samples. However, in sample C2, the proportion of lymphatic endothelial cells was significantly larger than its vascular counterpart. Sample C2 also had the highest percentage of T cells and contributed almost all the identified B cells. Due to this immune infiltration, to rule out mislabeling or patient misdiagnosis, we consulted with an experienced pediatric thyroid pathologist to re-evaluate the initial slides. Upon examination, the pathologist confirmed that immune recruitment was present and consistent with the histologic changes observed in response to the diagnostic fine-needle aspiration (FNA) performed prior to surgery. Even with the observed increased immune recruitment at the FNA injury site in sample C2, the proportion of cells identified as resident thyroid immune cells was substantially lower than previously reported in “normal” thyroid tissue collected from patients with thyroid malignancy (4, 8, 9). Spearman correlation analysis of the three samples based on their respective cluster proportions revealed high similarity between samples C1 and C3 (Figure 2D). Sample C2 had moderate correlation with the other samples, likely due to the already noted immune recruitment and increased proportion of T cells and B cells. Despite the differences in cluster proportions, correlation analysis of the clusters’ respective transcriptional profiles across the three samples confirmed their high transcriptional similarity and supported the inclusion of sample C2 (Figure 2E).

### Pathway analysis on nine main cell types

To determine pathways enriched in each cell type, we identified conserved markers across all samples and used them to perform gene set enrichment analysis (GSEA). Relevant biological pathways can be found in Figure 3. For thyrocytes (Thyrocyte 0 and 1 combined), 15 statistically significant GO pathways were retained. The most enriched pathways included those associated with iodide transport, thyroid hormone generation, metabolic processes, and pathways involved with ion transport (Figure 3A). These results are consistent with the known primary function of thyrocytes to produce thyroid hormone. Pathways involved with cellular polarization were also conserved across thyrocyte populations. Interestingly, immune pathways were found to be among the most downregulated pathways in thyrocytes, suggesting a potential immune-privileged or suppressive environment. To determine whether different thyrocytes may perform distinct functions within the follicular unit, conserved genes were identified within clusters Thyrocyte0 and Thyrocyte1, and used to perform pathway analysis. Within the Thyrocyte0 population, the only biological pathway with a positive normalized enrichment score was GOBP-G protein coupled receptor signaling pathway coupled to cyclic nucleotide second messenger. G protein coupled cyclic AMP signaling is downstream of the TSH receptor supporting activation of the TSH receptor in the Thyrocyte0 population. All remaining pathways had negative enrichment scores. This was in stark contrast to Thyrocyte1 in which many of the selected pathways had positive enrichment scores, including in multiple biosynthetic and metabolic processes. This data could be consistent with Thyrocyte0 responding more robustly to TSH, whereas Thyrocyte1 may be more metabolically active and utilizing energy for biosynthetic purposes. Thyrocyte populations were further analyzed and subclustered later in this study.

**Figure 3.**
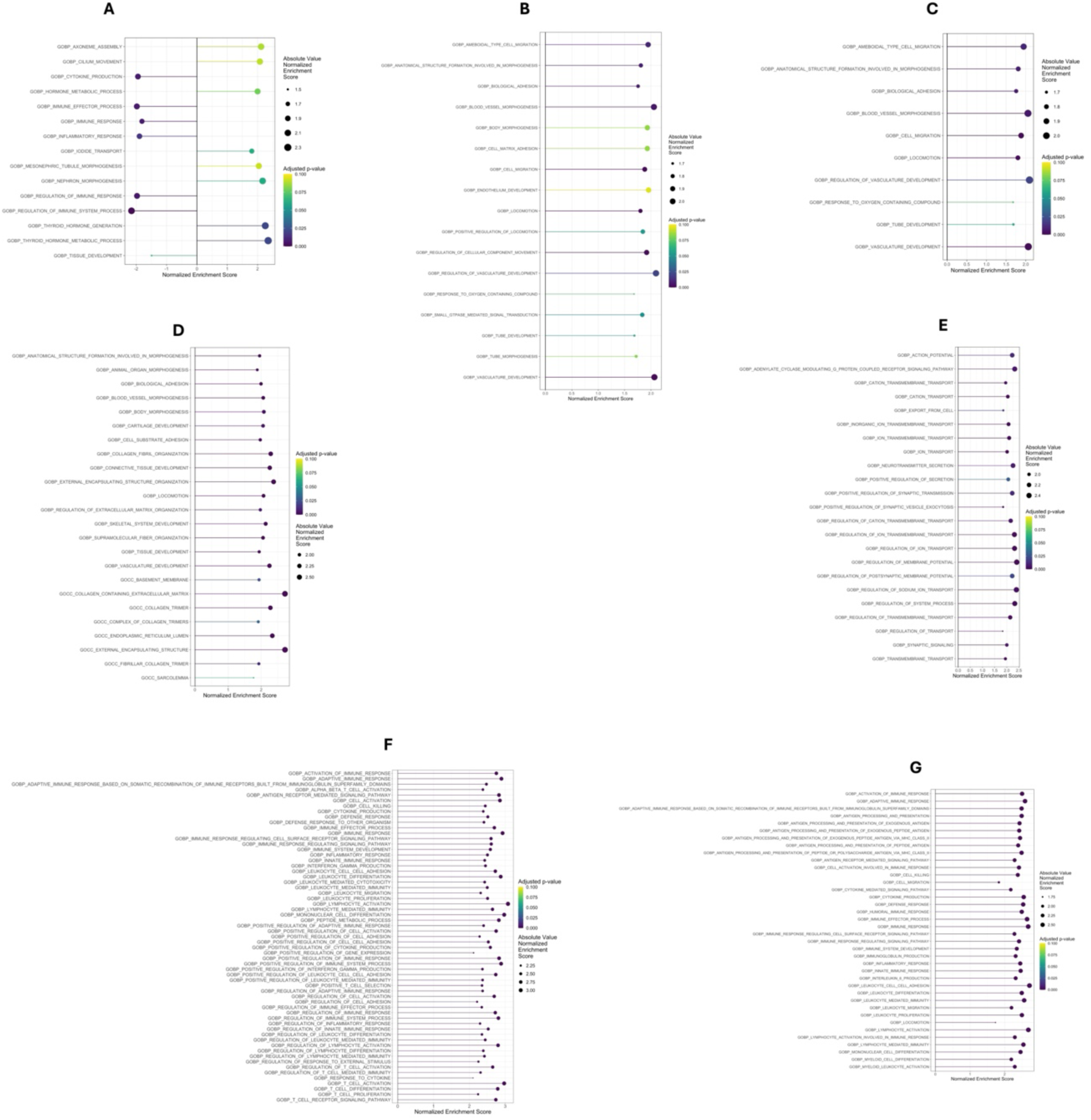
Selected enriched pathways by cell type/cluster. The results for each cell type were based on (ranked) genes within that specific cluster that were conserved across all three pediatric thyroid tissue samples and were filtered for entries having an adj. p-value ≤ 0.1. **(A)** Thyrocyte pathways, **(B)** Lymphatic endothelial cell pathways, **(C)** Vascular endothelial cell pathways, **(D)** Fibroblast pathways, **(E)** C cell pathways, **(F)** T cell pathways and **(G)** Myeloid pathways.

Seventeen GO pathways were selected for lymphatic endothelial cells (Figure 3B). By contrast, only 10 pathway entries were retained for vascular endothelial cells (Figure 3C). Pathways enriched in both groups of endothelial cells included processes required for the development of a robust vasculature and lymphatic drainage such as cell migration, locomotion, biological adhesion, and blood vessel morphogenesis.

Fibroblasts had 24 selected enriched pathways (Figure 3D). Analysis revealed upregulation of pathways involved in extracellular matrix organization, collagen synthesis and fibril organization, and morphogenesis. These pathways likely support the structural modifications that occur within the thyroid gland to organize thyrocytes into follicular units to produce thyroid hormone. Spatial transcriptomics and histological confirmation of fibroblasts and collagen deposition adjacent to follicular units will be necessary in future studies to investigate this hypothesis. We did not observe pathways associated with fibrosis, as has been reported in previous studies (4, 8, 9), nor did we observe any transcriptional signatures that would indicate activated TGF-β signaling in this population of pediatric fibroblasts.

Many pathways were identified among the rare cell populations, including C cells and immune populations. The increased pathway enrichment observed in these small clusters supports the unique and distinct identity of these cell types, despite their low abundance. Twenty-three pathways were found for C cells (Figure 3E), which included those involved in the hormonogenesis of calcitonin. Additionally, pathways involved in anion and cation transport were increased, supporting action potential generation, which is essential for hormone production. Pathways associated with “export from cells” were also increased, and support the secretion of calcitonin from these cells. The mixed T cell population returned 61 statistically significant pathways (Figure 3F), while the myeloid cluster had 38 pathways (Figure 3G). Within each immune population, pathways involved in activation of immune response and adaptive immune response were enriched. These data further support the presence of a resident immune population within the pediatric thyroid gland. No differentially expressed genes or enriched pathways were identified in the B cells, as this population was only attributed to sample C2.

### Thyrocyte subclustering

We were next interested in determining the heterogeneity of thyrocytes within the normal pediatric thyroid gland. Although sample C2 demonstrated high similarity to samples C1 and C3, we were concerned that the wound healing-associated immune infiltration and histologic changes may have altered thyrocyte biology. We therefore conservatively removed this sample from subcluster analysis. Thyrocyte cells (Thyrocyte0 and Thyrocyte1) from samples C1 and C3 were separated from the remaining clusters to create a new dataset. Using the SWANS pipeline and Clustree to avoid over-clustering, the cells subclustered into 7 distinct populations (Figure 4A). These subclustered thyrocytes were identified as Thyrocyte**<u>s</u>**0-6 to distinguish them from the initial two thyrocyte populations (Figure 4B). A breakdown of the number and proportion of cells in each new thyrocyte cluster is shown in Table 4. Both samples C1 and C3 contributed thyrocytes to each cluster. Interestingly, sample C3 had larger proportions of cells in clusters Thyrocytes1 and Thyrocytes2 compared to C1, whereas C1 had higher proportions of cells in clusters Thyrocytes3-6 compared to C3. No clusters were unique to a single sample, only distributed differently across the two samples.

**Figure 4.**
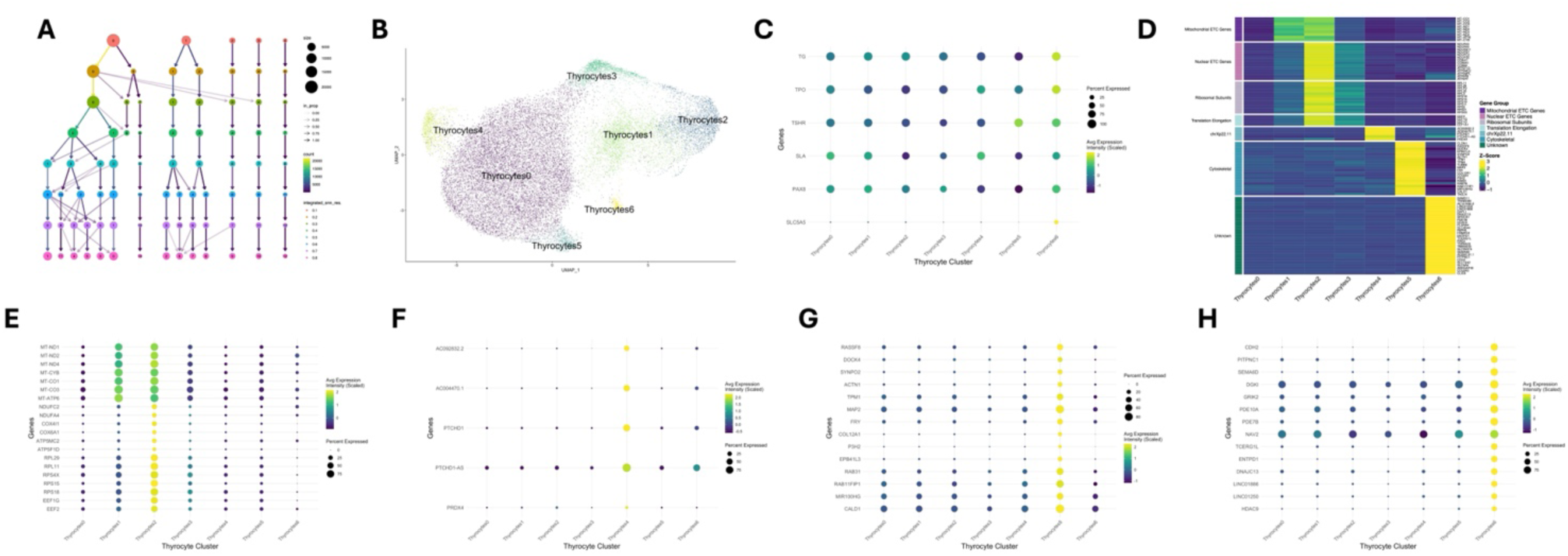
Single nuclei sequencing defines distinct, heterogeneous populations of thyrocytes within pediatric thyroid glands. All thyrocytes within samples C1 and C3 were subclustered to determine cellular heterogeneity specifically within the thyrocyte compartment. **(A)** Clustree image of thyrocyte cell-cluster assignment (and their potential reassignment) as a function of resolution. All cells are from samples C1 and C3. **(B)** UMAP plot of 23,093 thyrocyte cells from two pediatric thyroid glands (C1, C3) in seven thyrocyte subclusters. **(C)** Bubble plot of canonical thyroid marker genes *TG, TPO, TSHR, SLA, PAX8,* and *SLC5A5.* **(D)** Heatmap of selected cluster marker genes. Genes were selected and grouped based on their identified functional groups as discussed in the results. Gene expression was averaged across cells in each cluster and z-scores for each gene were calculated across subclusters. **(E)** Bubble plot of selected mitochondrial ETC, nuclear ETC, ribosomal subunit, and translation elongation genes identified as uniquely expressed in Thyrocytes1 and Thyrocytes2. **(F)** Bubble plot of all uniquely expressed genes in Thyrocytes4. **(G)** Bubble plot of selected cytoskeletal genes identified as uniquely expressed in Thyrocytes5. **(H)** Bubble plot of selected uniquely expressed genes in Thyrocytes6. For all bubble plots, average expression is depicted via dot color and the percentage of cells expressing the gene is denoted by dot size.

**Table 4:** Number of cells and their proportions in each of the 7 (Thyrocytes0-Thyrocytes6) thyrocyte subclusters subdivided by sample C1 and C3.

|  | **C1** | **C3** | **Total** |
| --- | --- | --- | --- |
| **Thyrocytes0** | 7790 (61.17) | 6322 (61.04) | 14112 (61.11) |
| **Thyrocytes1** | 1360 (10.68) | 1988 (19.19) | 3348 (14.5) |
| **Thyrocytes2** | 1240 (9.74) | 1165 (11.25) | 2405 (10.41) |
| **Thyrocytes3** | 939 (7.37) | 472 (4.56) | 1411 (6.11) |
| **Thyrocytes4** | 750 (5.89) | 219 (2.11) | 969 (4.2) |
| **Thyrocytes5** | 414 (3.25) | 186 (1.8) | 600 (2.6) |
| **Thyrocytes6** | 243 (1.91) | 5 (0.05) | 248 (1.07) |
| **Total** | 12736 (100.01) | 10357 (100) | 23093 (100) |

To explore the heterogeneity within the thyrocyte subclusters, we first evaluated the expression of canonical thyrocyte marker genes, including *TG, TPO, TSHR, SLA,* and *PAX8* (Figure 4C). We observed that a high percentage of cells across all thyrocyte subclusters expressed these genes, confirming their thyrocyte identity. Although the percentage of cells expressing these genes remained relatively consistent across all subclusters, the average expression within each subcluster varied significantly. Importantly, no single subcluster exhibited uniformly higher or lower expression across marker genes, indicating significant heterogeneity with no cluster-specific pattern. In contrast, *SLC5A5,* which is necessary for iodine transport into thyrocytes for thyroid hormone biosynthesis, was expressed in only a small fraction of cells across most subclusters (Figure 4C). Strikingly, Thyrocytes6 contained a much higher proportion of *SLC5A5-*expressing cells and exhibited significantly increased average expression compared to all other subclusters.

Given the clear heterogeneity between the thyrocyte subclusters, we next evaluated genes uniquely expressed within each subcluster. To identify these marker genes, we first averaged the expression of genes across cells within each subcluster and evaluated which were expressed at least three-fold higher in a given cluster compared to all others. The largest subcluster, Thyrocytes0, which comprised around 61% of all thyrocytes, did not uniquely express any marker genes. The lack of distinct markers suggests that these thyrocytes may represent a baseline or quiescent population from which the other subclusters diverge. Thyrocytes1, Thyrocytes2 and Thyrocytes3 all expressed a gene expression profile suggesting an energetically active state and engaged in protein synthesis. We collectively call these metabo-Thyrocytes. Thyrocytes1 and Thyrocytes2 uniquely expressed a set of genes encoded in the mitochondrial genome, including *MT-ND1, MT-ND2, MT-ND3, MT-ND4, MT-CYB, MT-CO1, MT-CO2, MT-CO3,* and *MT-ATP6* (Figure 4D). In addition to higher expression, the percentage of cells expressing these mitochondrial genes was also higher compared to the other clusters (Figure 4E). These genes encode core subunits of the electron transport chain (ETC) complexes, specifically NADH dehydrogenase (*MT-ND1, MT-ND2, MT-ND3, MT-ND4*), cytochrome c reductase (*MT-CYB*), cytochrome c oxidase (*MT-CO1, MT-CO2, MT-CO3*), and F-ATPase (*MT-ATP6*). High expression of these genes was specific to metabo-Thyrocytes, supporting that they represent unique, metabolically active sub-populations of thyrocytes. Interestingly, Thyrocytes2 was also significantly enriched for a distinct set of ETC-related genes encoded in the nuclear genome (Figure 4D). These gene products make up different subunits of the same complexes encoded by the mitochondrial genes upregulated in Thyrocytes1 and Thyrocytes2. Notably, these included subunits of NADH dehydrogenase *(NDUFA3, NDUFA4, NDUFB11, NDUFB7, NDUFC2, NDUFS5)*, cytochrome c oxidase *(COX4I1, COX6A1, COX8A)*, and F-ATPase *(ATP5F1D, ATP5MC2, ATP5MPL, ATP5PB, ATP5PF).* High expression of these nuclear-encoded ETC genes was specific to Thyrocytes2. However, moderate expression was also observed in Thyrocytes3 and, to a lesser extent, Thyrocytes1. Along with these ETC-related genes, Thyrocytes2 uniquely expressed genes encoding subunits of the ribosomal machinery *(RPL11, RPL29, RPLP2, RPL35, RPL3, RPS18, RPS15, RPS19, RPS11, RPS5, RPS21, RPS4X)* and involved in translation elongation (*EEF2, EEF1G, EEF1D, EEF1A1)*, both essential for protein synthesis (Figure 4D). These genes were also expressed in significantly more cells in Thyrocytes2 than any other cluster, indicating that in addition to being metabolically active, cells in Thyrocytes2 could be uniquely primed to support protein synthesis (Figure 4E).

Interestingly, Thyrocytes3 expressed the mitochondrial genes at lower levels and in fewer cells than Thyrocytes1 and Thyrocytes2. The nuclear ETC, ribosomal, and translation elongation genes were highest in Thyrocytes2, but still well expressed in Thyrocytes3, with higher levels observed than in Thyrocytes1. Thyrocytes3 also had moderately increased expression and percentage of cells expressing the mitochondrial ETC genes as compared to Thyrocytes0, Thyocytes4, and Thyrocytes5, but not nearly as robustly as observed in Thyrocytes 1 and Thyrocytes2 (Figures 4D, 4E). Taken together, these data support that Thyrocytes1-3 represent cells in dynamic flux or represent functionally distinct subsets of metabo-Thyrocytes.

Thyrocytes4 uniquely expressed only three genes, *PTCHD1, PTCHD1-AS,* and *PRDX4,* and two novel transcripts, AC092832.2 and AC004470.1 (Figure 4D). In addition to increased expression, a strikingly high percentage of cells in this cluster expressed these genes (Figure 4F). Interestingly, the three genes are all located on chrXp22.11 within 400 kb of each other and there are no other annotated genes within this region. The novel transcripts AC092832.2 and AC004470.1 directly overlap with *PTCHD1-AS* but are located on the opposite strand. These markers were highly expressed in cells from both samples C1 and C3, indicating their expression was not a sample-specific artifact. Additionally, *PTCHD1, PTCHD1-AS,* and *PRDX4* were all found to reside in the same topologically-associating domain in human cell line data, potentially explaining this co-expression (39). While these three genes and two novel transcripts were solely expressed in Thyrocytes4, their biological role in this population remains unclear.

Cells within the Thyrocytes5 subcluster were specifically enriched for a set of cytoskeletal-related genes, including those involved in maintenance of cell-cell junctions *(CLDN1, RASSF8, DOCK4)*, actin dynamics (*SYNPO2*, *ACTN1, TPM1, TPM4, TAGLN),* microtubule dynamics (*TUBB6, MAP2, FRY)*, collagen chain assembly *(COL12A1, COL8A1, P3H2)*, cytoskeletal-membrane linkage (*EPB41L3*), and vesicle transport (*RAB31, RAB7B, RAB11FIP1)* (Figure 4D). We called these cytoskel-Thyrocytes. These genes were expressed at least three-fold higher and in a much higher percentage of cells in this subclusters compared to others (Figure 4G). Interestingly, we also found significant upregulation of *MIR100HG* (3.1-fold compared to the next highest expressing cluster), a lncRNA that positively regulates *CALD1* expression by targeting miR-142-5p (40). *CALD1* is an actin- and myosin-binding protein that stabilizes the actin filament network and was expressed 3.25-fold higher in cytoskel-Thyrocytes compared to the next highest expressing cluster. Upregulation of these cytoskeletal genes was not seen in any other subcluster, indicating cytoskel-Thyrocytes likely represent a distinct structural population or are uniquely primed for intracellular movement of thyroid hormone via vesicle transport.

Thyrocytes6 contained the fewest cells of any subcluster, accounting for only around 1% of thyrocytes. Cells in this cluster were almost exclusively from sample C1 (Table 4). Many genes were uniquely expressed in this cluster; however, pathway analysis did not reveal any biologically relevant functions. Notably, many of the genes upregulated in this cluster are classically linked to different aspects of the central nervous system and are highly expressed in neuronal tissue *(SEMA6D, NAV2, CDH2, GRIK2, PDE10A)* (Figures 4D, 4H). Interestingly, *SLC5A5,* the gene encoding the sodium-iodide symporter, was also highly expressed in this cluster (10.8-fold compared to the next highest expressing cluster). However, due to its atypical composition and extremely small size, we did not attempt further characterize this thyrocyte subcluster or infer its biological role.

To confirm that low viability was not the driver of thyrocyte subclustering, particularly for Thyrocytes4 and Thyrocytes6, we examined the proportion of transcripts mapping to the mitochondrial genome. For every cluster, the percent of mitochondrial reads was less than 15% (Figure S2). While we observed fluctuations in the percent of mitochondrial reads across subclusters, they displayed a similar pattern across samples C1 and C3, additionally confirming that sample-specific artifacts were not driving thyrocyte subclustering. We next evaluated cell cycling within each subcluster to determine whether the distribution of cells across cell cycle phases was abnormal. We found that each cluster contained cells in various phases of the cell cycle and that cells in each phase did not uniquely originate from either sample C1 or C3 (Figure S3). This further confirms that thyrocyte subclustering is driven by distinct transcriptional profiles and not by abnormal cell cycle states or decreased viability.

To determine whether the observed gene expression differences influenced functional pathway activation, we performed GSEA. GSEA was performed using the fgsea package after z-transforming expression values and generating per-cluster ranked gene lists. The number of significant (adjusted p-value ≤ 0.1) pathways ranged between 81-185 per cluster. Eighteen pathways were enriched in all clusters and included pathways related to thyroid hormone biogenesis (Table 5). These data support the idea that even though multiple thyrocyte populations were identified through subclustering, all thyrocytes maintain gene expression programs to produce thyroid hormone and polarize into follicular units. Shared pathways also included signal processing and responses to stress. Interestingly, in all thyrocyte populations except Thyrocytes2, the most negatively enriched pathways were involved in immune recognition or chemokine signaling, supporting an immune suppressive role for these thyrocyte populations.

**Table 5:** Pathways (18) present in all seven (Thyrocytes0-6) clusters.

|  |
| --- |
| **Shared Pathways** |
| SA FAS SIGNALING |
| HP LARGE POSTERIOR FONTANELLE |
| SIG PIP3 SIGNALING IN B LYMPHOCYTES |
| GOBP REGULATION OF CELLULAR COMPONENT MOVEMENT |
| GOBP THYROID HORMONE GENERATION |
| GOBP MYOBLAST PROLIFERATION |
| SA PTEN PATHWAY |
| SIG INSULIN RECEPTOR PATHWAY IN CARDIAC MYOCYTES |
| SIG IL4RECEPTOR IN B LYPHOCYTES |
| GOBP REGULATION OF MYOBLAST PROLIFERATION |
| GOMF OLFACTORY RECEPTOR ACTIVITY |
| HP ABNORMAL CIRCULATING THYROGLOBULIN LEVEL |
| SIG BCR SIGNALING PATHWAY |
| NABA SECRETED FACTORS |
| GOBP CELLULAR RESPONSE TO STRESS |
| SIG PIP3 SIGNALING IN CARDIAC MYOCTES |
| SIG CHEMOTAXIS |
| HP DELAYED PROXIMAL FEMORAL EPIPHYSEAL OSSIFICATION |

In addition to shared pathways, we also sought to identify pathways that were unique to each thyrocyte population. The most dominant cluster, Thyrocytes0, was the only cluster enriched for the Stem Cell Division pathway, and could therefore represent a progenitor population. Metabo-Thyrocytes (Thyrocytes1 and 2) had enrichment for pathways related to energy production. NADH Dehydrogenase Complex, Electron Transport Chain, and Respiratory Chain Complex IV-associated pathways were found in Thyrocytes1, whereas Oxidative Phosphorylation and Respiratory Electron Transport Chain pathways were unique to Thyrocytes2. Thyrocytes2 also had pathways with increased ribosome activity and processes involved with protein trafficking and transport, such as Protein Folding in ER and Cytoplasmic side of the lysosomal membrane. These results are consistent with the enrichment of ribosomal genes and translation elongation genes in Thyrocytes2 (Figures 4D, 4E). This suggests that although the two subpopulations share expression of many of the same genes, their functional roles may differ and potentially represent distinct stages of energy production within the electron transport chain. Despite sharing similar gene expression with the other metabo-Thyrocytes, there were no unique pathways enriched in Thyrocytes3 related to thyroid function. Although the uniquely expressed genes in Thyrocytes4 did not indicate a distinct functional role of this subpopulation, the pathways unique to this population indicate a population actively responding to stress. Thyrocytes5, noted above to be cytoskel-Thyrocytes, showed unique enrichment in pathways related to phospholipase activity. Upon closer examination, the genes represented in these pathways were also found to participate in cytoskeletal remodeling and in responses to extracellular cues that may induce cytoskeletal reorganization. Thyrocytes6 returned the most unique pathways, consistent with the large number of uniquely expressed genes. However, this cluster contained only 1% of the identified thyroid follicular cells, and we are not confident this represents biologically relevant variability in function of this population.

The expression of well-known thyroid differentiation scoring (TDS) genes (as defined by the Thyroid Cancer Genome Atlas project (41)) were plotted to evaluate the expression of each TDS gene in all the cells of each thyrocyte cluster to determine if they varied across the 7 thyrocyte populations (Figure 5A-P). Cluster Thyrocytes3 had on average, the lowest expression of most of the thyroid-specific genes. Interestingly, all the clusters displayed heterogeneity in expression of *TSHR, PAX8, GLIS3, NKX2-1, SLC26A4, THRB, DIO1, DIO2,* and *SLC5A8* whereby numerous cells within each of these clusters displayed extraordinarily little to no expression of these genes. Even though *SLC5A5* is necessary for iodine transport into thyrocytes and thyroid hormone biosynthesis, *SLC5A5* expression was extremely low in all thyrocyte cells, regardless of cluster. As all patients had normal TSH or free T4 levels, this demonstrates that low basal expression of *SLC5A5* and therefore NIS is sufficient to support thyroid hormone production (Table 1). Pendrin (*SLC26A4*) expression increased from Thyrocytes0 to Thyrocytes2, whereas *GLIS3* followed the opposite trend and expression decreased from Thyrocytes0 to Thyrocytes2. GLIS3 directly regulates expression of pendrin (42), so it was quite unusual that their expression patterns are opposite. GLIS3 also regulates the expression of *TG,* which was found to be higher in Thyrocytes2 than in Thyrocytes0. This supports the idea that the expression of the proteins involved in hormone biosynthesis may have a more complex regulatory expression system than originally proposed. *TG, TSHR, PAX8, GLIS3, TPO, SLC26A4*, and to a slightly lesser degree, *NKX2-1, DIO1, DIO2*, and *SLC5A8* were expressed in most cells and in most clusters. *DUOX1* and *DUOX2* were only expressed in clusters Thyrocytes1 and Thyrocytes2, and *THRA* and *SLC5A5* were not detected in most cells, regardless of cluster (Figure 5A-P).

**Figure 5.**
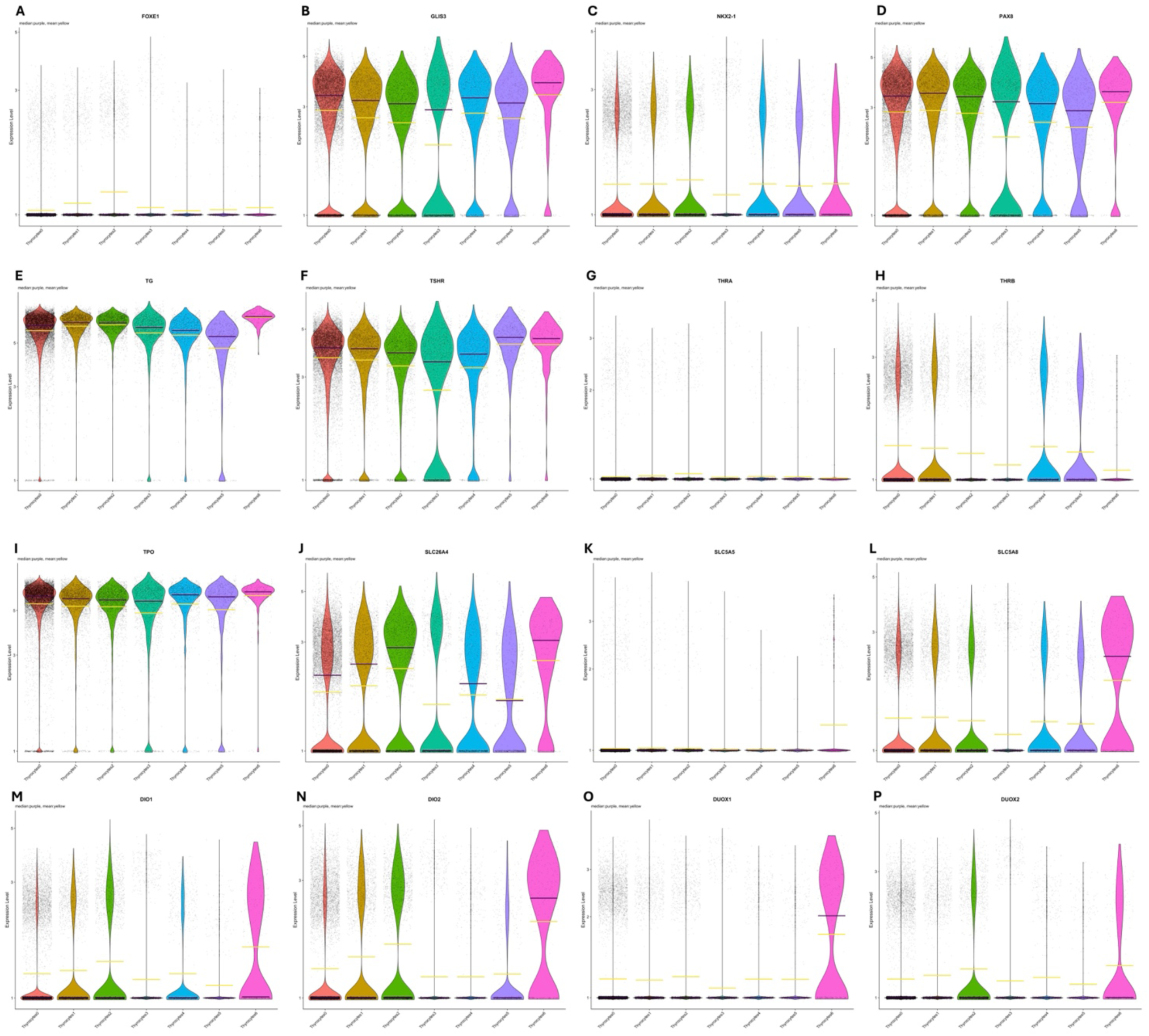
Violin plots of normalized log expression of thyroid specific genes. Expression of thyroid differentiation genes were determined. Each dot represents the expression of the given gene in each cell within the indicated clusters. The yellow line indicates the mean and the purple line indicates the median. **(A)** *FOXE1* **(B)** *GLIS3* **(C)** *NKX2-1* **(D)** *PAX8* **(E)** *Thyroglobulin* (*TG*) **(F)** Thyroid Stimulating Hormone Receptor (*TSHR*) **(G)** *Thyroid Hormone Receptor α* (*THRA*) **(H)** *Thyroid Hormone Receptor* β (*THRB*) **(I)** *Thyroid Peroxidase* (*TPO*) **(J)** *Pendrin* (*SLC26A4*) **(K)** *Sodium Iodide Symporter*, (*SLC5A5*) **(L)** *SLC5A8* **(M)** *DIO1* **(N)** *DIO2* **(O)** *DUOX1* **(P)** *DUOX2*.

## DISCUSSION

To analyze thyroid snRNA-seq data, we utilized SWANS, an automated pipeline to analyze scnRNA-seq data and compare resulting clustering arrangements in an interactive environment. The pipeline included typical scnRNA-seq workflow operations: data cleaning, dimensionality reduction and imputation, clustering, and differential gene expression (DGE) analysis/gene set enrichment analysis (GSEA).

Our analysis included tissue from three pediatric thyroid samples. Two distinct endothelial cell clusters were identified and classified into two groups: vascular and lymphatic endothelial cells. These identities were further supported when looking at each population’s DEGs and associated pathways. Cellular adhesion, blood vessel morphogenesis, cell migration, and cell locomotion were common pathways found in both groups of endothelial cells. The classical lymphatic endothelial marker *LYVE1* was expressed in the lymphatic group, but not in the vascular endothelial cells. Supporting the wound healing observed in sample C2 in response to a prior FNA, we observed increased lymphatic endothelial cells and a greater proportion of lymphatic to vascular endothelial cells in this sample. It is possible there are other subgroups of endothelial cells, and that heterogeneity exists even within these two subclusters. However, because of the relatively low number of cells in each group (1087 lymphatic endothelial cells and 345 vascular endothelial cells), subclustering these populations was deemed to not be biological relevant or informative.

Fibroblasts were found in all samples in almost equal abundance, supporting the presence of a resident population of fibroblasts within the normal thyroid. Differential gene expression analysis revealed the most upregulated genes within the fibroblast population included *LAMA2, SLIT2, COL3A1,* and *COL5A2*. Upregulated pathways involved in extracellular matrix organization, collagen synthesis, fibril organization, and morphogenesis were observed. Enrichment of these pathways supports fibroblasts’ role in the development of the thyroid gland and its structure. The abundance of fibroblasts in this study is much less than what has been reported in previous studies of “normal” thyroid tissue (4, 8, 9). It is possible these differences are due to the age of the patients from which samples were collected, although Hong et al. (4) did not report significant differences in the cellular composition between young (20-30 years old) and old (50-60 years old) patients. We hypothesize that “normal” thyroid tissue adjacent to malignant tumors is not truly normal and that both transcriptional and cellular recruitment changes occurred in these adjacent areas, even in the absence of overt histological changes. Further, the phenotype of fibroblasts adjacent to malignant tissue supports a pathological role, as these cells expressed genes associated with fibrosis and pathological TGF-β signaling (4). Future studies will need to directly compare normal thyroid tissue obtained from samples with no malignant transformation within the gland to adjacent normal areas in thyroids with malignant tumors. Our results and predicated cellular composition of the thyroid gland more closely resemble what has been reported in the zebrafish thyroid, and support that one should be cautious of assuming tissue adjacent to a thyroid tumor is directly comparable to normal thyroid tissue in the absence of malignancy.

A mixed population of T cells and myeloid cells were observed in all three pediatric thyroid samples. The highest percentage of T cells and myeloid cells were found in sample C2, corresponding with evidence of a healing FNA site, although these cells were still in very low abundance (3.86% and 0.61%, respectively). Similar to what we reported with fibroblasts, our samples’ combined “resident” immune population was significantly smaller than the immune populations reported in other studies that also profiled “normal” thyroid tissue adjacent to thyroid tumors (4, 8, 9). These studies reported that up to 50% of cells in these “normal” thyroids were of immune origin. It is possible that some of these variations could also be related to sample processing, as our samples were flash frozen, whereas previous studies have utilized both fresh and frozen samples for individual cell profiling. B cells were found almost exclusively in sample C2 and represented 1.32% of the sample’s total cells. The presence of T cells and myeloid cells in all three pediatric samples support a resident immune population within the thyroid. Although we did not observe increased expression of markers of regulatory T cells, the population of identifiable T cells was too small for robust subclustering and analysis. Pathways involved in the activation of innate and adaptive immune responses were enriched within each immune population. Immune suppression is also supported by the finding that Thyrocytes0 and Thyrocytes1 had pathways enriched for immune suppression and regulation. Future studies with a larger number of samples and profiled cells can more comprehensively define the predicted crosstalk between thyrocytes and stromal cells within the normal thyroid gland.

Comparing our data to those of studies profiling “normal” thyroid adjacent to malignancy supports that both cellular recruitment to the thyroid and the transcriptional profiles of thyroid cells can be altered throughout the gland when malignant transformation is present. It will be critical for future studies to include more samples to increase the number of profiled immune cells. This will help determine whether the resident immune cells impart an immune suppressive phenotype in the normal thyroid during non-pathogenic conditions. Additionally, identifying neutrophils in scRNA-seq datasets requires special and timely tissue handling and processing (43). When analyzing the data, devising a way to distinguish neutrophils from discarded low-feature cells will be needed to capture this cell type. We expect to see resident and recruited neutrophils in normal and malignant thyroid (fresh) tissue; however, neutrophils do not survive tissue freezing and would not be available in our dataset. Future thyroid single-cell studies looking to describe the full immune cell repertoire should consider the neutrophils’ sensitivity in their initial study design. Including neutrophils in future analyses will be critical, as tumor-infiltrating neutrophils have been associated with poor prognosis in many tumors (44), although their role in thyroid tumorigenesis is underexplored.

C cells were the least abundant population of cells found in the dataset and were only identified in samples C1 and C2 by high expression of *CALCA, NRXN1, NKAIN2, CALCB, MEG3*, and *STK32A*. However, this is the first study to our knowledge that has reported the presence of C cells in the thyroid through scnRNA-seq. This is not surprising as C cells are reported to represent only about 2-4% of the cells within the thyroid (45). Here, we report that C cells comprise 0.04% of the cells identified in our thyroid tissue samples. The smaller proportion identified in this study could be due to variability in C cell abundance across different ages or the decreased stability of these cells during freezing. It is also possible that our samples did not contain sufficient lateral portions of the thyroid, where C cell concentrations are the highest.

Our initial analysis to identify cell types present in the pediatric thyroid categorized two thyrocyte populations: Thyrocyte0 and Thyrocyte1. These two populations were identified across the three-sample dataset and contained 66% and ∼ 26% of the cells, respectively, and represented the most abundant cell type. This is stark contrast to a recent report by Hong et al. that reported the presence of only 35% thyroid epithelial cells in normal young and old thyroid (4). The *SLA* gene was one of the most differentially expressed genes when thyrocytes were compared to all other cell types, and was used along with *TPO, TG, PAX8, TSHR, NKX2-1, SLC5A5* and *SLC26A4* for annotation purposes. Unfortunately, snRNA-seq is not ideal for detecting low-level transcripts, and this is likely why these thyroid-specific genes did not initially reach the threshold for inclusion as markers. *RMST* was significantly overexpressed in thyrocytes compared to all other cell types in the samples, and its expression was found in 97.8% of thyrocytes, compared to only 22.3% of all other cells. *RMST* is a long noncoding RNA that has been reported to be overexpressed in the thyroid compared to other tissue, and has recently been implicated in differentiation of thyroid cells (46). De Martino et al. report that *RMST* is downregulated in anaplastic thyroid cancer (ATC) and that cell growth and invasiveness of ATC cells can be inhibited by re-expression of *RMST*. Up- and downregulation of many different lncRNAs have been reported in thyroid cancer; however the role that these genes play in normal thyroid cell physiology is underexplored. Expression of such genes in normal thyrocytes would implicate some role in normal physiology in addition to pathologic phenotypes. Interestingly, many lncRNAs were uniquely expressed in distinct thyrocyte subclusters, particularly in Thyrocytes4, Thyrocytes5, and Thyrocytes6. While almost all of these genes lacked functional annotations, our findings further underscore the importance of lncRNAs in normal thyroid function and suggest that they may play distinct roles in different cellular sub-populations. Future studies should investigate the biological relevance of these lncRNAs, as well as how their differential expression may shape cellular phenotypes.

We went on to further dissect the transcriptional heterogeneity of thyrocytes by performing a subclustering analysis solely on thyrocytes. This yielded seven distinct populations of thyrocytes showing vast heterogeneity within these subclusters. While all clusters expressed canonical thyrocyte markers, the expression levels and percent of cells expressing these genes varied significantly with no clear pattern across all subclusters. To further interrogate the biological relevance of these different subclusters, we identified marker genes that were uniquely expressed in one or several subclusters compared to others. We identified no uniquely upregulated or downregulated marker genes for Thyrocytes0, the largest population, suggesting a possible quiescent or baseline thyrocyte population from which the other subclusters differ. We further identified three metabolically active sub-populations which we called metabo-Thyrocytes: Thyrocytes1, Thyrocytes2, and Thyrocytes3. While all three subclusters had increased expression of genes involved in oxidative phosphorylation and protein synthesis, the patterns of expression varied significantly. We identified Thyrocytes2 as displaying the highest expression and percentage of cells expressing mitochondrial ETC genes, nuclear ETC genes, ribosomal subunit genes, and translation elongation genes, suggesting this might be the most metabolically active subpopulation. Interestingly, Thyrocytes1 had comparable expression of mitochondrial ETC genes, but reduced expression of nuclear ETC genes and protein synthesis-related genes compared to Thyrocytes2, supporting a potential shift in activation or function. Thyrocytes3, while having the lowest mitochondrial ETC gene expression of the three subclusters, had higher expression of nuclear ETC genes, ribosomal subunit genes, and translation elongation genes than Thyrocytes2. Due to this unique heterogeneity, we hypothesize that these three metabolically active subclusters may represent thyrocytes in different phases of activation. It is possible that in response to TSH, thyrocytes activate a specific gene expression program to begin the synthesis of thyroid hormone. Since thyroid hormone synthesis is an energy-intensive process, upregulation of genes involved in ATP synthesis is expected. Because expression of mitochondrial genes can be more rapidly induced than nuclear genes, it is possible that they are the first to be upregulated in this gene expression program. As the time after stimulation increases, ETC genes in the nuclear genome become upregulated to generate protein subunits necessary to generate ATP. At the same time, genes involved in protein synthesis also become upregulated to generate the machinery needed to synthesize thyroid hormone. As these transcripts are synthesized into mature proteins and the cells reach the required protein levels, induction of these genes is reduced, and their RNA expression levels fall. Future studies with spatial transcriptomics and proteomic profiling are necessary to test this hypothesis.

Furthermore, since snRNA-seq captures cells at a fixed point in time, it is possible that we are capturing cells at various timepoints after stimulation and thus in various phases of this gene expression program. Future spatial transcriptomic studies could seek to determine whether thyrocytes from these different subclusters reside adjacent to each other, or whether entire follicles may be synchronized to a specific sub-population of thyrocytes. Regardless of whether these different transcriptional signatures represent distinct cell populations of distinct fates, or different states of thyrocyte activation, these data support a heterogenous population of thyrocytes and represent more diversity within thyrocyte cell populations than previously reported.

We additionally identified many cytoskeletal genes as markers of Thyrocytes5, cytoskel-Thyrocytes. These genes were involved with basic cytoskeletal organization, as well as intracellular transport. Due to the unique upregulation of these genes, we hypothesize that cells in cytoskel-Thyrocytes may be responsible for thyroid hormone trafficking from the colloid to endothelial cells and into circulation. In order for thyroid hormone to be released into circulation, it must be endocytosed and packaged into vesicles at the apical membrane of follicular thyrocytes, transported through the cell, and exocytosed at the basolateral membrane. Within a cell, vesicular transport relies on both the presence of organized cytoskeletal filaments and signaling proteins, usually from the Rab GTPase family, which modulate vesicle formation and transport. Cytoskel-Thyrocytes uniquely express proteins controlling the assembly of the two major filaments involved in vesicular transport (actin and microtubules). Additionally, cytoskel-Thyrocytes uniquely express many Rab GTPase family genes, which are necessary for targeted vesicle transport. Unique expression of genes involved in the vesicular transport system, as well as other cytoskeletal-stabilizing genes, suggest that these thyrocytes may play a role in thyroid hormone transport out of the follicular lumen.

While we identified these unique thyrocyte sub-populations and postulated their potential biological functions, it remains unclear whether these subclusters reflect stable cell types or transient cell states. It is possible that each thyrocyte has a unique biological function and retains this function for its entire lifespan, such that each subcluster represents a unique and persistent cell type. Alternatively, all thyrocytes may be capable of cycling through these functions, with each subcluster reflecting a particular phase captured at the time of snRNA-seq. If the latter is true, it also remains unclear whether thyrocytes within a given follicle transition through these in a coordinated manner, such that every cell is in the same state at the same time, or whether individual cells progress asynchronously.

Each of the thyrocyte subclusters had conserved enrichment of pathways related to thyroid hormone biosynthesis, supporting the hypothesis that all thyrocytes maintain the ability to produce hormones. However, pathway analysis revealed distinct activity within each subcluster. Some clusters were more metabolically active, whereas others had increased expression of ribosomal genes and pathways associated with hormone trafficking through the cell for exportation into the blood stream. Quite surprising was the variability of *DIO2* expression across thyrocyte clusters. *DIO2* encodes the gene for the type II iodothyronine deiodinase enzyme, which converts T4 to T3. This differential expression suggests that only a subset of thyrocytes within the thyroid may be responsible for thyroid hormone conversion to the active form T3. Future studies will be needed to determine whether this local conversion of T4 to T3 is for secretion of active T3 into circulation or for local action of thyroid hormone within the gland. *DUOX2* was expressed at the highest level in the cluster Thyrocytes2. *DUOX2* is responsible for H_2_O_2_ production and is used by *TPO* in the biosynthesis of thyroid hormone (Figure 1A). *TPO* expression was elevated across all populations of thyrocytes, but the selective increase of *DUOX2* in cluster Thyrocytes2 suggests that hormone production occurs selectively in distinct thyrocyte subpopulations. Collectively, the subclustering analysis supports the concept that although all thyrocytes have the ability to produce hormones, different steps in the hormone biogenesis pathway may be preferentially performed in certain thyrocytes. Unfortunately, snRNA-seq is not ideal for detection and quantification of low-level transcripts within cells. It is likely that the true variability within thyrocytes populations may be even greater than is detectable through snRNA-seq. As the data presented represents a single snapshot in time, the heterogeneity within thyrocytes may represent dynamic changes in gene expression that do not represent true cell type differences, but instead, dynamic cell states that are in continual flux.

Apart from the B cell and C cell populations, we did not observe any additional bias in the number of cells originating from an individual sample. These results suggest that there are indeed common lineages of thyrocytes and that these cell types are consistent across samples. For this analysis, we limited the cohort to the female sex to obviate potential confounding influences as we know thyroid maladies are associated with different incident rates depending on sex. Additionally, we maintained a closely-matched age range in the patient population to avoid observing transcriptional differences that could be attributed to age-related developmental and hormonal changes. These studies lay the foundation for future work, and include markers, cell types, and their proportions to confirm their identification in other cohorts across sex and age ranges. Future studies will benefit from additional samples, inclusion of patients from different ages, and inclusion of both male and female patients.

Significantly, to our knowledge, this is the first study to define cellular populations and thyrocyte heterogeneity in the normal pediatric thyroid, or any human thyroid completely absent of malignant transformation. These data suggest that “normal” thyrocytes and adjacent stromal cells in the background of malignancy are likely not truly normal, as the percentages and gene expression profiles of these cells are different than thyroid tissue in the absence of malignancy. Ideally, future studies will be able to profile tissue from patients with no evidence of thyroid changes, but collection of this tissue is extremely difficult and limited. This analysis and the work of Gillotay (3) support the heterogenous nature of thyrocytes. However, it is still not clear how each of these different thyrocytes are arranged in the follicular structure and how these thyrocytes coordinate thyroid hormone biogenesis and export of mature thyroid hormone into circulation. We observed some variations in the cellular composition of thyroids from different patients which may be due to sampling bias, as the exact location that cells were extracted from was not the same from each patient. Two samples were from the right lobe, whereas one sample was from the left lobe. There may be under-appreciated differences in cellular composition within the lobes of the thyroid gland or isthmus. Our future studies will seek to validate these findings through spatial transcriptomics and histological investigations to define the location of these cells within the thyroid.

## DATA AVAILABILITY

The data presented in this study are available from the corresponding author (ATF) upon reasonable request.

## ETHICS STATEMENT

This study involved human-derived biospecimens and was reviewed by the Children’s Hospital of Philadelphia Institutional Review Board. The study was approved (IRB #20-018240) for waiver of consent as these were existing biospecimens from previous surgeries.

## AUTHOR CONTRIBUTIONS

ERR, JCRF, and ATF designed the study. ERR designed and developed bioinformatic tools for data analysis. KH, AI, TB, AJB and ATF identified and assessed clinical features of the patient cohort. JCRF and ZS performed the experimentation and acquired the data. ERR, NEB, JCRF, TB and ATF analyzed and interpreted the results. ERR, NEB, JCRF, AJB, and ATF drafted and edited the manuscript. All authors contributed to the article and approved the submitted version.

## ACKNOWLEDGEMENTS

We thank the entire Franco Laboratory for thoughtful discussion and critique while performing this study and writing this manuscript. We thank Dr. Mei Zhang, Xiaoxu Yang, and Aliza Yousef in the Children’s Hospital of Philadelphia Single Cell Technology Core, and Dr. Teodora Orendovici and Stephen Mahoney from the High Throughput Sequencing Core.

