## Supplementary_Figures for "Single-cell analysis reveals the cellular and transcriptional diversity of thyrocytes in the normal pediatric thyroid"

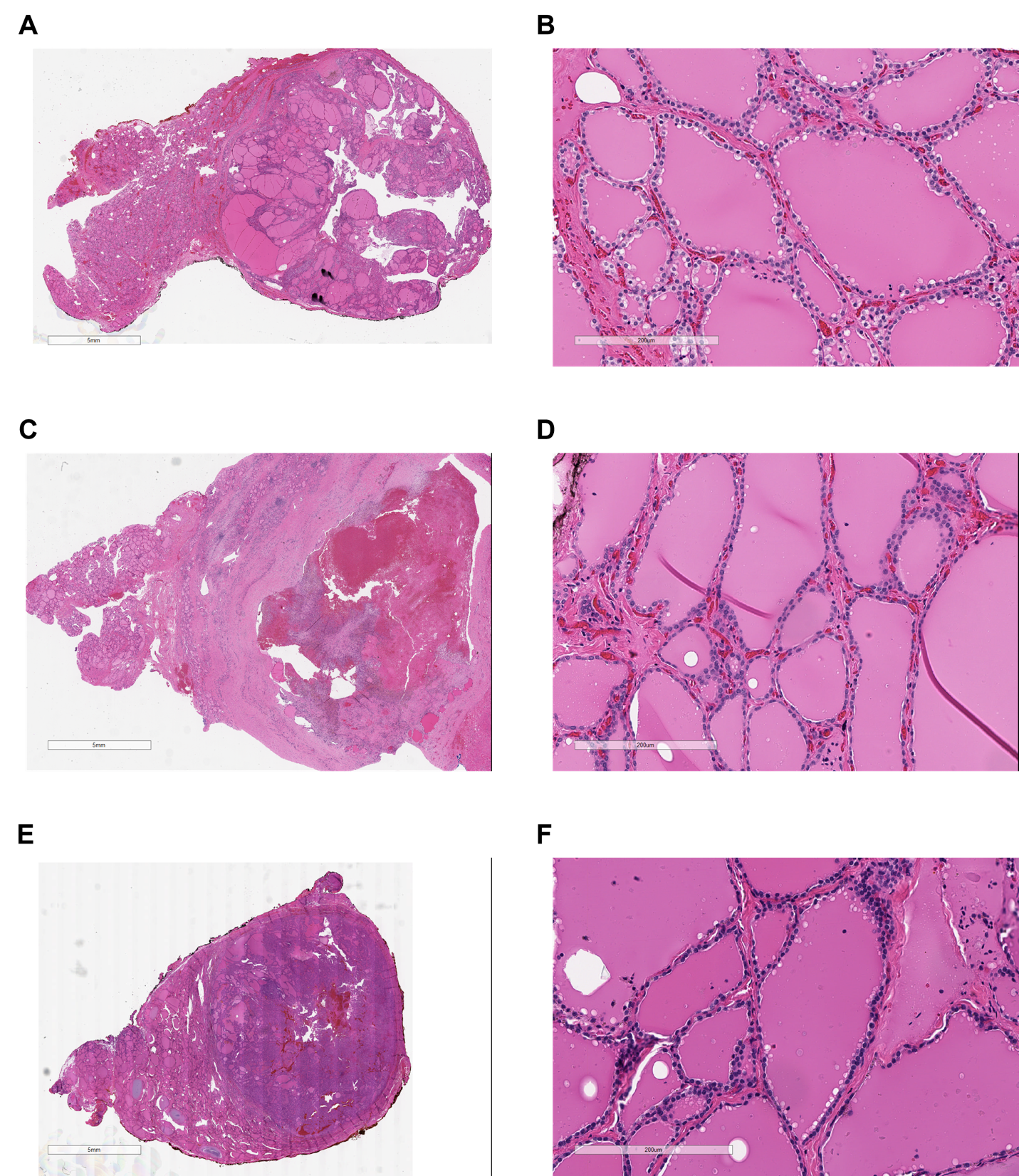

**Figure S1.** Histology of samples for snRNA sequencing. **(A-F)** Representative H&E-stained sections of sample C1 at low magnification **(A)** and higher magnification **(B)**, sample C2 at low magnification **(C)** and higher magnification **(D)**, and sample C3 at low magnification **(E)** and higher magnification **(F)**. White scale bars on low magnification images represent 5mm, black scale bars on higher magnification images represent 200µm.

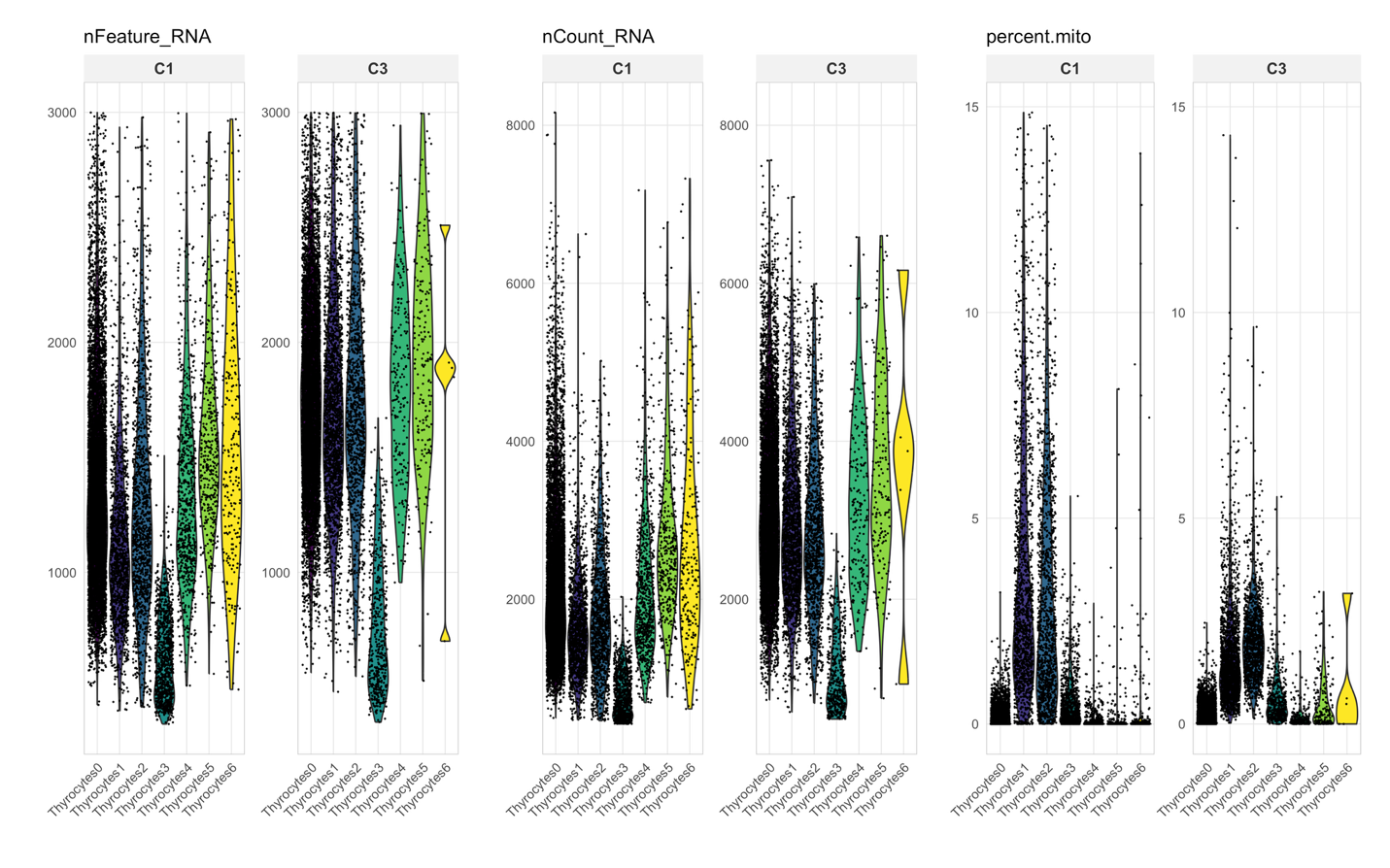

**A**

**B**

**C**

**Figure S2.** Violin plots of number of genes, number of molecules, and mitochondrial transcript percentage for each thyrocyte sub-subcluster. **(A)** Violin plots displaying the number of genes detected in each cell for each sub-cluster. **(B)** Violin plot displaying the number of distinct RNA molecules in each cell for each sub-cluster. **(C)** Violin plots displaying the percentage of mitochondrial transcripts in each cell for each sub-cluster. Each dot represents a unique cell and cells are stratified based on sub-cluster assignment.

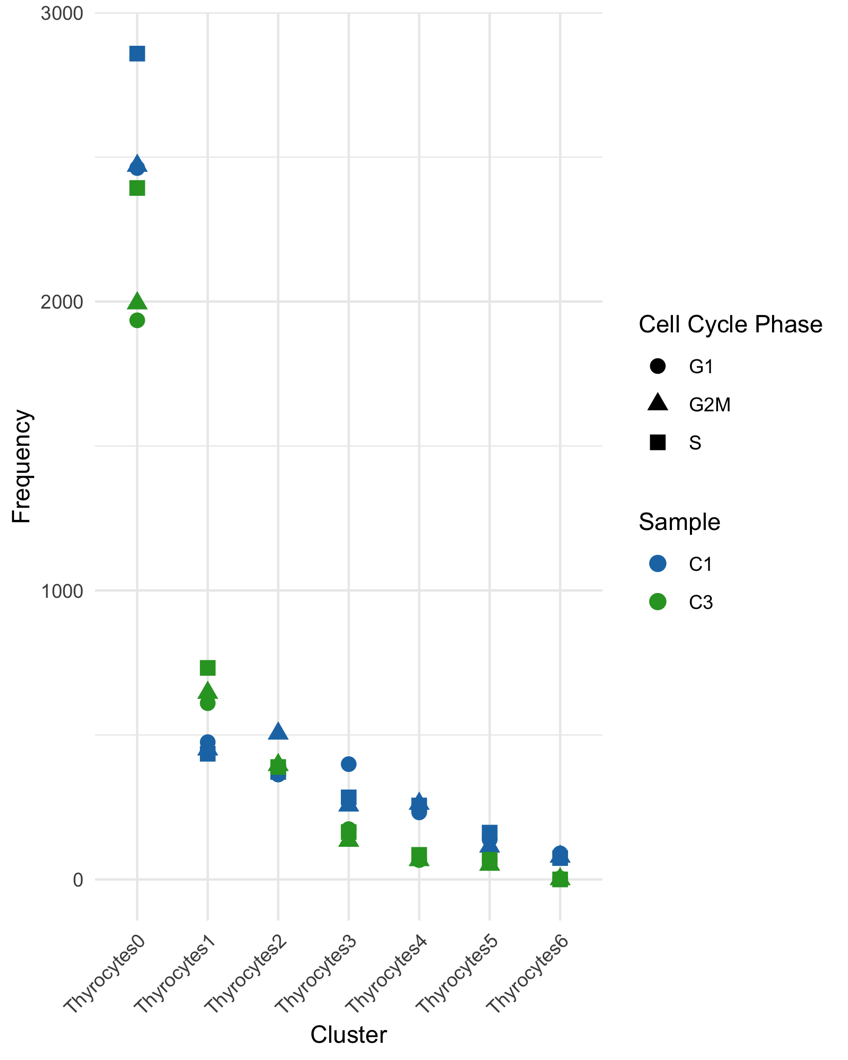

**Figure S3.** Plot showing cell cycle phase of cells in each thyrocyte subcluster. Cell cycle phase is denoted by dot shape and colored based on which sample the cells originated.
